# Population structure of *Phytophthora infestans* collected from potatoes in Ecuador, Colombia, Peru, Bolivia and Uruguay

**DOI:** 10.64898/2026.01.14.699522

**Authors:** Myriam Izarra, Mario Coca-Morante, Pérez Willmer, Laura Sánchez, Soledad Gamboa, Diana Valle, Viviana Cuarán, Beatriz Guerra-Sierra, Jan Kreuze

**Author notes:** Correspondence author: Beatriz Guerra, Jan Kreuze.

## Abstract

Late blight, a destructive disease affecting potatoes, is caused by the oomycete *Phytophthora infestans* and remains a major threat to potato production worldwide. Understanding the population structure of this pathogen is essential for effective disease management. We examined the genetic structure of South American *P. infestans* populations using 182 isolates: 97 from Bolivia and southern Peru, 14 from Colombia, 57 from Ecuador (1993-2022), and 14 from Uruguay. The isolates were characterized by clonal lineage, mitochondrial haplotype, and mating type using multilocus genotypes based on microsatellite (SSR) markers. In Bolivia, only the lineage/haplotype 2A1/Ia was identified, whereas Peru exhibited both 2A1/Ia and EC1/IIa, all of which were mating type A1. Puno was the sole department where both lineages were present. In historical Ecuadorian populations, we found US1/Ib and EC1/IIa (*P. infestans*), as well as EC2/Ic and EC3/Ia (*Phytophthora andina*), while recent populations showed only US1. In Colombia, EC1/IIa and a distinct clonal lineage (CO4/Ia) were identified within the present dataset. In Uruguay, 2A1/Ia was predominant.

These results provide updated insights into the genetic diversity and geographic distribution of *P. infestans* across South America and highlight the importance of continued regional surveillance for improved late blight management.

## Introduction

*Phytophthora infestans*, the causal agent of potato late blight, has a complex population structure and evolutionary history in South America. The global migration history of *P. infestans* has been the subject of extensive research and debate. There are various hypotheses regarding the origin of *P. infestans*, including Mexican, Andean, and hybrid origin hypotheses. Goss et al. (2014) suggested that Mexico was the center of origin from which the pathogen migrated to South America, then to the United States, and finally to Europe (Andrivon, 1996). A study by Gómez-Alpizar et al. (2007) found that Andean populations harbored greater nucleotide and haplotype diversity than Mexican populations, and that the oldest mutations in *P. infestans* originated in South America, supporting the Andean origin theory. More recently, Coomber et al. (2025) reaffirmed that the Andes is the center of origin, and Patarroyo et al. (2024) suggested a Peruvian origin with a probability of 0.631. The movement of genotypes contributes to shaping population structure and disease dynamics (Guha Roy et al., 2021).

The study of the population structure of *P. infestans* is complex and dynamic and has been widely studied across different regions. It is influenced by factors, such as natural migration or migration associated with the movement of infected potato seed tubers, sexual reproduction, and geographical isolation. Additionally, *P. infestans* can rapidly evolve and adapt, posing a challenge for disease management (Cooke & Lees, 2004; Dey *et al*., 2018; Ludwiczewska *et al*., 2025). Microsatellite markers, also known as simple sequence repeats (SSRs), are highly polymorphic co-dominant nuclear markers, providing valuable insights into the genetic variation and population dynamics of this devastating pathogen (Lees *et al*., 2006).

*P. infestans* is potentially able to reproduce sexually when compatible A1 and A2 mating types coexist in the same population. However, sexual reproduction is not always observed even when both mating types are present, as compatibility between strains may vary (Runno-Paurson *et al*., 2012). Until the 1980s, only mating type A1 was known to occur in Europe, which restricted the pathogen to asexual reproduction. The appearance of A2, first detected in Switzerland and likely introduced through potato imports from Mexico (Rhouma *et al*., 2024), changed the disease dynamics (Drenth et al., 1993; Shattock, 2002). The presence of both A1 and A2 mating types allows sexual reproduction, potentially leading to increased genetic diversity and the formation of oospores that, as alternative inoculum sources, can persist in the soil and act as long-term infection sources (Drenth *et al*., 1993; Shattock, 2002; Yuen & Andersson, 2013). This supplementary source of inoculum, combined with increased genetic diversity, enhances the pathogen’s capacity to adapt and persist in the absence of a host, potentially complicating traditional disease management strategies (Drenth et al., 1993, 1994).

Clonal lineages of *Phytophthora infestans* are designated using standard nomenclature (e.g., US1, EC1, 2A1), while mitochondrial haplotypes are indicated by Roman numerals (e.g., Ia, IIa). Combined designations (e.g., US1/Ib, EC1/IIa, 2A1/Ia) refer to the association between clonal lineage and mitochondrial haplotype. In Peru, the population structure of *P. infestans* exhibits clonal lineages with varying dominance across regions and over time. In the southern region, isolates collected between 1994 and 1999 were characterized by clonal lineages and mitochondrial haplotypes. In Cusco, the identified lineages were US1/Ib, EC1/IIa, PE3/Ia, PE5/IIa, and PE6/IIa. In Puno, US1/Ib, PE3/Ia, and EC1/IIa were found (Perez *et al*., 2001).

A subsequent study analyzing isolates collected between 1997 and 2013 reported the presence of PE3/Ia and EC1/IIa in Cusco, US1/Ib and PE3/Ia in Puno, and EC1/IIa in Apurímac (Lindqvist□Kreuze *et al*., 2020). More recently, isolates from 2017 were analyzed by Lindqvist□Kreuze et al. (2020), revealing EC1/IIa in Cusco and both EC1/IIa and PE7.1/Ia (a variant of PE7) in Puno, all of which belonged to the A1 mating type.

*P. infestans* populations in Bolivia were studied using herbarium specimens to determine clonal lineages and mitochondrial haplotypes, which were found to be US1/Ib, and from modern isolates, which identified BR1/IIa (Saville & Ristaino, 2021). Gabriel et al. (2018) identified the A1 mating type, in contrast to Goodwin, S.B. et al. (1994), who identified mating type A2 in Bolivia.

The population structure of *P. infestans* in Colombia has been reported to be dominated by the clonal lineage EC1/IIa, whereas CO1/IIa and CO2/IIa have also been identified on potato (Vargas *et al*., 2009). In contrast, US1/Ib and CO3/Ib were identified on tomato hosts (Céspedes et al., 2013; Gilchrist Ramelli et al., 2009; Vargas et al., 2009; Olave-Achury et al., 2022).

In Uruguay, the presence of BR1/IIa was previously reported (Deahl *et al*., 2003), but more recent studies have revealed the presence of 2A1/Ia (F. Lucca et al., 2023).

In Ecuador, the presence of distinct clonal lineages and mitochondrial haplotypes has been studied over the years, with EC1/IIa being dominant on potato hosts and US1/Ib on tomato hosts (Forbes *et al*., 1997; Oyarzun Pj *et al*., 1998). In addition to *P. infestans*, closely related species such as *Phytophthora andina* have also been reported in the Andean region. *P. andina* is considered a distinct species within the *P. infestans* complex, likely of hybrid origin, and is associated with infections on wild and cultivated *Solanum* species (Gómez-Alpizar *et al*., 2008; Oliva *et al*., 2010). Additionally, lineages such as EC2 and EC3 have previously been identified as *P. andina* on different Solanum species (Gómez-Alpizar *et al*., 2008).

The importance of studying South American *P. infestans* populations lies in their considerable diversity (Coomber *et al*., 2025), which has implications for disease management and breeding resistance. Clonal lineage 2A1, first reported in Europe in the early 1980s (Cooke *et al*., 2012), was dominant on potatoes before 2006 (Mariette *et al*., 2016). The 2A1 lineage has also shown significant expansion beyond Europe including the eastern-Africa region (EAR), comprising Kenya, Uganda, Tanzania, Burundi, and Rwanda (Njoroge *et al*., 2019). Its presence in South America has been associated with the introduction of infected potato tubers. This lineage is now regarded as the dominant one in Argentina, Chile, Brazil, and Uruguay (Acuña et al., 2019; F. Lucca et al., 2023)

We propose that the genetic variation and population dynamics of *Phytophthora infestans* in South America are influenced by both geographic and agroecological factors, leading to distinct regional patterns. In particular, we anticipate that populations in ecologically linked or neighboring regions (such as the central Andes—Colombia, Ecuador, Peru and Bolivia) will exhibit higher genetic similarity, while those geographically distant from this region (such as southern Uruguay) will show more pronounced genetic differences possibly influenced by the importation of seed potatoes from Northern Hemisphere countries.

## Materials and methods

### Sampling

A total of 182 *Phytophthora infestans* isolates collected between 1993 and 2025 from five South American countries (Ecuador, Colombia, Peru, Bolivia and Uruguay) were included in this study. Of these, 127 isolates were from field collections conducted between 2017 and 2025 in Bolivia (n=67), Peru (n=30), Colombia (n=14), Uruguay (n=14), and Ecuador (n=2, collected in 2022). Additionally, 55 historical Ecuadorian isolates collected between 1993 and 2008 were obtained from the CIP *Phytophthora* collection and had been preserved in liquid nitrogen.

During 2017-2019, leaf samples with late blight-like symptoms were collected in Bolivia and southern Peru. Bolivian samples originated from the department of Cochabamba, whereas Peruvian samples were collected in the departments of Cusco, Apurimac, Ayacucho, and Puno. In Colombia, samples were collected in 2022 from Santander and the North of Santander. Two Ecuadorian samples were collected in 2022 from Pichincha. In Uruguay, samples were collected between 2023-2025 from Canelones and San Jose. Each leaf sample yielded a single isolate for further molecular characterization.

Field samples (Bolivia, Peru, Colombia, Uruguay, and Ecuador-2022) were collected on FTA cards (Whatman FTA Classic Card, catalog number WB120055; GE Healthcare UK Ltd.) according to the manufacturer’s guidelines (Cooke, 2020). Each sample was placed on FTA paper with the sporulating side down and individually crushed in the field (Gamboa *et al*., 2019). The FTA cards were then stored in zip-lock bags and shipped to the pathology laboratory at the International Potato Center (CIP) in Lima, Peru, for processing.

Complete metadata for all isolates, including collection year and date, geographic origin (country, department, province, and district), altitude, host, and genotypic characterization, are provided in Supplementary Table S1.

### Genotypic Analysis

All 182 *Phytophthora infestans* isolates were subjected to molecular characterization. Mitochondrial haplotypes (mtDNA) and multilocus genotypes (MLGs) were determined for all isolates using PCR-based assays and a 12-plex SSR panel as described below.

Field-collected samples were obtained on FTA cards as described above. The 55 historical Ecuadorian isolates (1993-2008), preserved in liquid nitrogen at the CIP collection, were reactivated in culture and transferred onto FTA cards prior to molecular analysis.

DNA extracted from FTA cards (Qiagen^®^) was performed following the manufacturer’s instructions with minor modifications: two 3 mm discs were excised (instead of one), rinsed twice in modified Tris (hydroxymethyl) aminomethane-ethylenediaminetetraacetic acid TE^-1^ buffer (10 mM Tris and 0.1 mM EDTA), and incubated in 50 μl TE buffer (10mM Tris and 1mM EDTA), at 95 °C for 5 min, and stored at −20 °C until use.

### Mating Type

Mating type information was available for 178 of 182 isolates included in this study. For field-collected isolates, mating type was determined using W16 CAPS assay (Judelson *et al*., 1995). For the historical Ecuadorian isolates (1993-2008), mating type information was obtained from previous in vitro pairing assays recorded in the CIP database.

#### W16 CAPS assay

The mating type (A1 and A2) was determined using PCR primers W16-1 (5’-AAC ACG CAC AGG CAT ATA AAT GTA-3’) and W16-2 (5’-GCG TAA TGT AGC GTA ACA GCT CTC-3’), as described by Judelson et al. (1995) for all isolates except the Ecuadorian isolates. PCR reactions (20 μL) contained 4 μL of 5x buffer (Gotaq, Promega), 6 μL of MgCl_2_ (25 mM), 0.2 μL of dNTPs (100 μM), 0.4 μM of each primer, 1 U of Taq DNA polymerase (Gotaq^®^, Promega), and 10 ng of template DNA. The thermal cycling protocol included an initial denaturation at 94 °C for 5 min, followed by 30 cycles of denaturation at 94 °C for 1 min, primer annealing at 53 °C for 1 min, and elongation at 72 °C for 1 min, with a final elongation step at 72 °C for 10 minutes. Afterwards, the PCR products were digested for 3h at 37 °C with Hae III (New England Biolabs, #R0108) in a 20 μL reaction containing 10 μL of PCR product, 2 μL of CutSmart Buffer 10X, 0.3 μL enzyme, and nuclease-free water (Brylińska *et al*., 2018). Digested fragments were separated on a 2% agarose gel in TBE buffer at 40 V for 4 h and visualized under UV light using GelRed (Biotium^®^) as nucleic acid stain.

#### Pairing Assay

For the historical Ecuadorian isolates, mating type had been previously determined by pairing with A1 (*P. infestans*) and A2 (*P. andina*) tester strains using dual-cultures on 10% clarified V8 agar. Plates were incubated at 15-18 °C in the dark and examined microscopically for oospore formation at the hyphal interfaces within the contact zone. Isolates forming oospores with A1 and A2 testers were classified as A2 and A1, respectively.

### Mitochondrial haplotype

Mitochondrial haplotypes were determined for all 182 isolates using a PCR-restriction fragment length polymorphism (PCR-RFLP) assay following Griffith & Shaw (1998). A specific mitochondrial region was amplified using primer P2, and PCR products were digested with MspI.

Restriction fragments were separated on a 2% agarose gel in TBE buffer at 40 V for 4h and visualized under UV light using GelRed (Biotium^®^) as nucleic acid stain. Haplotypes were assigned by comparing with reference isolates representing the established Ia, Ib, IIa, and Ic mitochondrial haplotypes (Griffith & Shaw, 1998).

### Molecular Markers

All 182 isolates were genotyped using a 12-plex microsatellite (SSR) panel following a modified protocol of Li et al. (2013) and Saville et al. (2016). PCR amplification was performed using the Type-It Microsatellite PCR kit (QIAGEN) according to Saville et al. (2016).

Amplified products were submitted to Arizona State University for each sample, 2 μL of a 1:50 diluted PCR product was combined with formamide containing LIZ500 size standard ladder and loaded onto 3730 instrument. Alleles sizes were determined using GENEMARKER v1.9 (SoftGenetics) according to the allele binning parameters described by Li et al. (2013). Clonal lineage assignment was performed by comparison with reference SSR profiles representing previously genotyped clonal lineages, including 2A1, EC1, BR1, PE3, PE7, PE7.1, and US8 (Vargas *et al*., 2009; Martin *et al*., 2019; Lindqvist□Kreuze *et al*., 2020; Saville & Ristaino, 2021). Additional Andean reference lineages included EC2, EC3, and PE8, previously reported within the *P*.*andina* (Adler *et al*., 2004; Oliva *et al*., 2010; Forbes *et al*., 2016) which were associated with the Andean species *P*.*andina*.

### Data Analysis

All statistical analyses were conducted using R version 4.3.1 (R Core Team, 2019). The following packages were used: polysat (v.1.7.7), poppr (v.2.9.7), RColorBrewer (v.1.1-3), magrittr (v.2.0.3), ape (v.5.8-1), and ggtree (v.3.12-0). Locus and population-level statistics, including the number of multilocus genotypes (MLGs), were calculated using the package poppr (Kamvar *et al*., 2014).

Clonality was assessed using the index of association (Ia) and the standardized index of association (r□d), with statistical significance evaluated through 999 permutations using the ia() function in poppr. For diversity analyses sensitive to repeated multilocus genotypes (MLGs), clone correction was applied by retaining a single representative of each MLG per population.

Population structure was explored using principal coordinates analysis (PCoA), hierarchical clustering, and minimum spanning networks (MSN) all based on Bruvo’s genetic distance (Bruvo *et al*., 2004). For PCoA, a Bruvo distance matrix was calculated for triploid SSR genotypes using the meandistance.matrix function in the R package polysat (Clark & Jasieniuk, 2011), and ordination was performed using classical multidimensional scaling (cmdscale) in R. Hierarchical relationships were examined using a unweighted pair group method with arithmetic mean (UPGMA) dendogram generated from a Bruvo distance matrix calculated with bruvo.dist (specifying locus-specific repeat lengths) and clustered using upgma in poppr. Genotypic relationships were further visualized using a minimum spanning network (MSN) constructed with bruvo.msn in □poppr (Kamvar *et al*., 2014). Ordination and network analyses were performed using the full dataset to reflect genotype frequencies. Reference genotypes representing known clonal lineages were included across analyses to support lineage assignment.

Diversity statistics were computed for each population, including the number of individuals (N), number of multilocus genotypes (MLG), expected number of multilocus genotypes (eMLG), standard error (SE), Shannon-Wiener Index (H), Stoddart and Taylor’s index (G), Simpson’s index (λ), evenness (E.5), and expected heterozygosity (Hexp), index of association (Ia), and standardized index of association (r□D). Additional locus-level statistics included allelic richness, Simpson’s diversity index (1–D), expected heterozygosity (Hexp), and evenness (Shannon, 1948, 2001; Simpson, 1949; Hedrick, 1999).

Analysis of molecular variance (AMOVA) was performed using Bruvo’s genetic distance to partition genetic variation among clonal lineages and among geographic locations nested within clonal lineages (∼ lineage/location). AMOVA was conducted on a clone-corrected dataset, retaining one representative per multilocus genotype (MLG) per population prior to distance matrix computation. The analysis was implemented using the poppr.amova function with the ade4 method. Statistical significance was assessed using 999 Monte Carlo permutations.

## Results

### Samples collected

A total of 182 *Phytophthora infestans* isolates were analyzed in this study. In Bolivia, 67 samples were collected from the department of Cochabamba, across ten provinces. In Peru, samples were obtained from four departments: Cusco (n=1), Apurímac (n=4), Ayacucho (n=6), and Puno (n=19). In Colombia, 14 samples were collected from Norte de Santander (n=6) and Santander (n=8). In Ecuador, 57 isolates were obtained from ten departments: Carchi (n=11), Bolívar (n=2), Chimborazo (n=2), Cotopaxi (n=5), Imbabura (n=3), Morona Santiago (n=1), Napo (n=9), Pastaza (n=2), Tungurahua (n=8), and Pichincha (n=14). Two isolates from Pichincha were collected in 2022, while the remaining Ecuadorian isolates correspond to historical samples preserved in the CIP collection. In Uruguay, 14 samples were collected from the departments of Canelones (n=10) and San José (n=4). Detailed isolate metadata are provided in Tables 1 and S1.

**Table 1.**
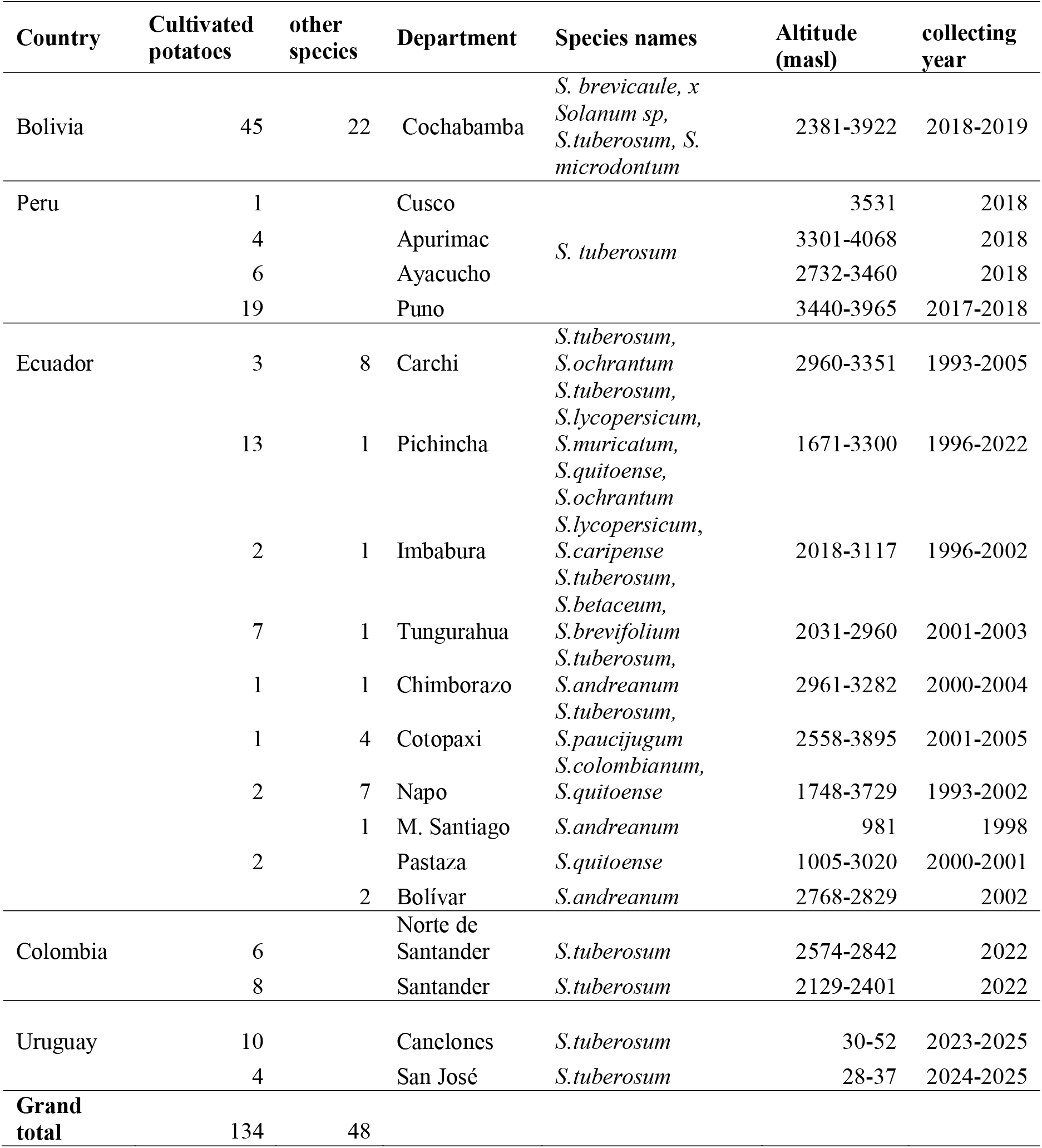
Summary of samples collected and analyzed in this study.

### Mating type test

The primer pair W16–1 and W16–2 amplified a 557-bp fragment in all isolates analyzed with this assay. Following digestion with HaeIII, A1 isolates were identified by the presence of the undigested 557-bp fragment, together with the expected digested fragment (457-bp), consistent with the expected banding pattern for A1 mating type.

All isolates from Bolivia, Peru, Colombia, and Uruguay were classified as A1 based on the W16 assay (Figure S1, Table S1). For Ecuadorian isolates, mating type was determined using pairing assays. Most isolates were A1, while a single isolate (Ec-3365), obtained from the wild species *Solanum brevifolium*, was identified as A2 and corresponded to *Phytophthora andina* (Table S1).

### Genotypic characterization

#### Mitochondrial haplotypes

Analysis of the mitochondrial haplotypes showed that isolates from Bolivia (n=67), Uruguay (n=14), Peru (n=8), Colombia (n=10), and Ecuador (n=2) belonged to haplotype Ia. In contrast, IIa was detected in 23 isolates from Peru, 3 from Colombia, and 38 from Ecuador. Haplotype Ib was detected in 16 Ecuadorian isolates, including two collected in 2022. A single isolate (Ec-3365) obtained from the wild species *Solanum brevifolium* corresponded to haplotype Ic (Figure S2, Table S1).

#### Population structure

Principal coordinates analysis (PCoA) based on Bruvo genetic distance revealed clustering patterns among isolates from 19 departments across five countries, together with nine reference genotypes. The analysis showed that the reference genotype PE7.1 clustered with the globally distributed clonal lineage 2A1 (Figure 1).

**Figure 1.**
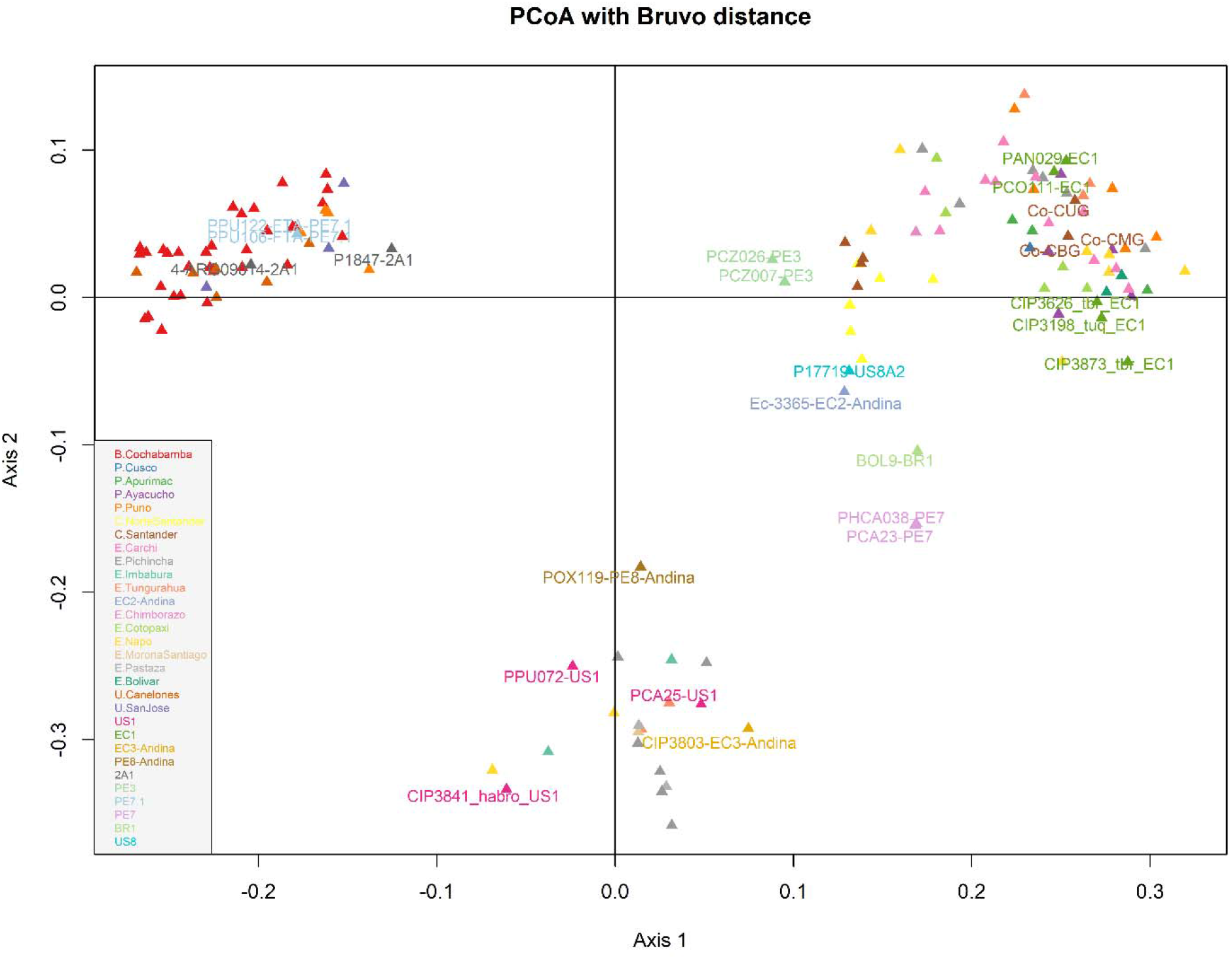
Principal coordinate analysis (PCoA) based on Bruvo’s genetic distance calculated from SSR marker data for *Phytophthora infestans* and *P*.*andina* isolates. Colors represent geographic origin: Bolivia (Cochabamba, red), Peru (Cusco, blue; Apurimac, green; Ayacucho, purple; Puno, orange), Colombia (Norte de Santander, bright yellow; Santander, brown), Ecuador (Carchi, pink; Pichincha, gray; Imbabura, teal; Tungurahua, coral; Chimborazo, light fuchsia; Cotopaxi, lime green; Napo, soft yellow; MoronaSantiago,sandy beige; Pastaza, light gray; Bolivar, forest green); and Uruguay (Canelones, burnt orange; San José, indigo). Reference genotypes used for clonal lineage assignment are indicated: US1(fuchsia), EC1(olive green), 2A1 (dark Gray), PE3 (mint green), PE7.1 (sky blue), PE7 (lavender) and BR1 (pale green), US8 (turquoise) for *P. infestans*; and EC2 (dusty blue), EC3(mustard), PE8 (bronze) for *P. Andina*.

Hierarchical clustering using a UPGMA dendrogram based on Bruvo’s distance identified six major clonal lineages (2A1, US1, EC1, EC2, EC3, and a distinct clonal lineage, here designated as CO4, identified in Colombia based on its clustering pattern relative to reference genotypes included in the dataset) (Figure 2).

**Figure 2.**
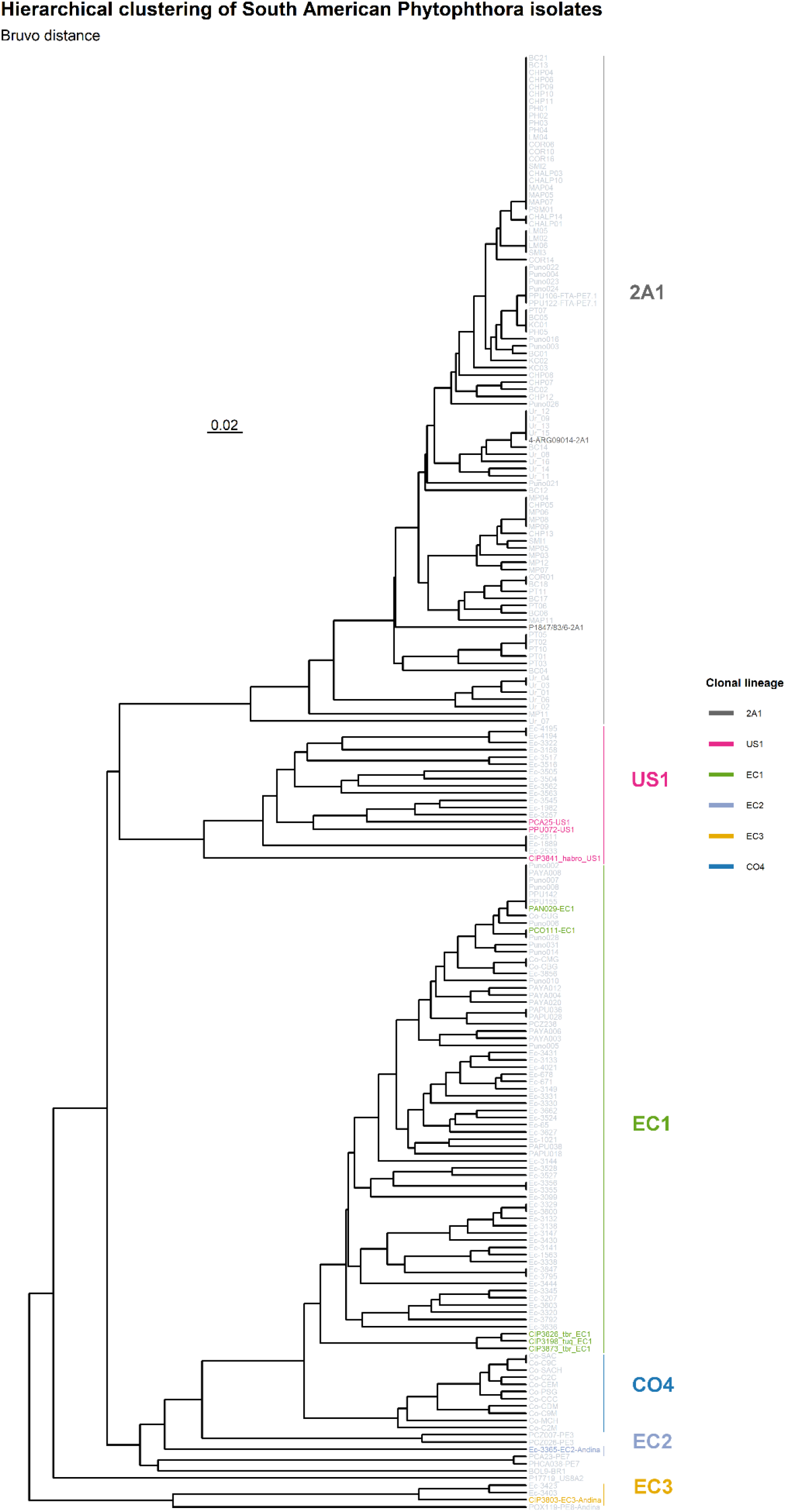
UPGMA dendrogram of *Phytophthora* isolates from Bolivia, Peru, Colombia, Ecuador, and Uruguay based on Bruvo’s distance from 12 SSR loci. Clonal lineage assignment was inferred from clustering with included reference genotypes. Major lineages (US1, EC1, 2A1, EC2, EC3, and CO4) are indicated; EC2 and EC3 correspond to *P. andina*, whereas the remaining lineages belong to *P. infestans*.

#### Clonal lineage distribution

In Bolivia (n=67) and Uruguay (n=14), all isolates belonged to the 2A1 clonal lineage. In Peru, isolates from Puno were distributed between EC1 (n=11) and 2A1 (n=8). In contrast, isolates from Cusco (n=1), Apurimac (n=4), and Ayacucho (n=6) were all assigned to EC1, indicating that this lineage predominated in southern Peru (73.3%), followed by 2A1 (26.6%).

In Ecuador, US1 was not detected in cultivated potato but occurred in several host species, including *S. lycopersicum* (18.8%), *S. muricatum* (12.5%), *S. andreanum* (6.2%), *S. quitoense* (56.3%) and *S. caripense* (6.2%). In contrast, EC1 was detected in cultivated potatoes (39.5%) and in several wild *Solanum* species, including *S. colombianum* (18.4%), *S. ochrantum* (23.7%), *S. paucijugum* (10.5%), and *S. andreanum* (7.9%).

Within the Ecuadorian population, EC1 represented 66.7% of isolates, followed by US1 (28.1%) and EC3 (3.5%, all of which were associated with *S. betaceum*). EC2 represented 1.8% of isolates and was detected in *S. brevifolium*. The largest number of isolates were collected in Carchi and Pichincha. In Carchi, all isolates belonged to EC1, whereas in Pichincha the population consisted of EC1 (42.9%) and US1 (57.1%). The isolates collected in 2022 from Pichincha belonged to the US1 lineage and represented 14.3% of the isolates from that department. In Tungurahua, three clonal lineages were detected: EC1 (62.5%), EC2 (12.5%), and EC3 (25%).

In Colombia, two well-differentiated clonal lineages were detected. The newly identified lineage CO4 represented 78.6% of isolates and occurred in both studied departments and 21.4% corresponded to EC1, found only in Santander (Figure 2, 3).

**Figure 3.**
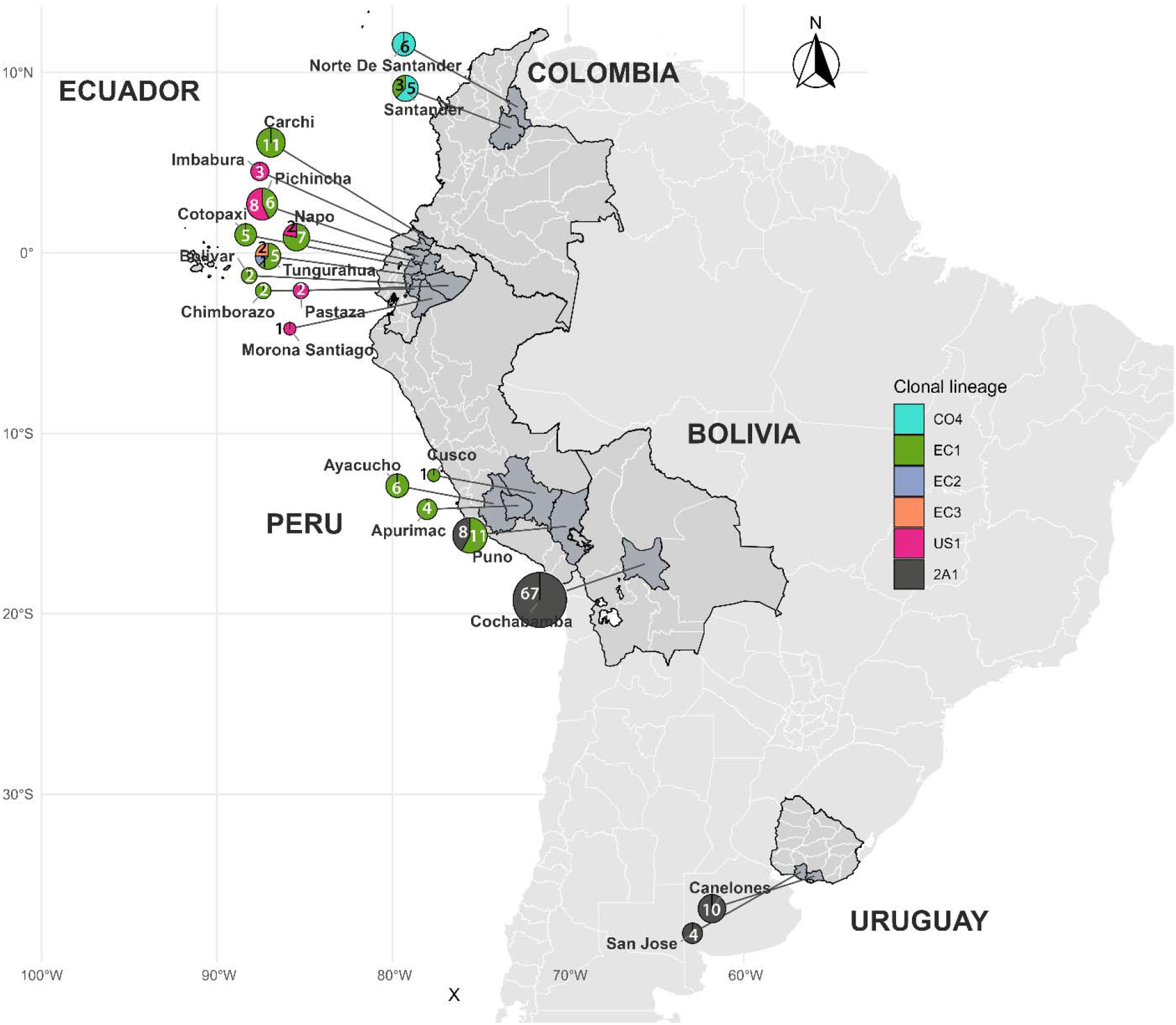
Geographical distribution of *Phytophthora* isolates (n =182) collected in Bolivia and Peru (2017–2019), Ecuador (1993–2022), Colombia (2022), and Uruguay (2023–2025). Departments highlighted in gray indicate sampled locations, and pie charts show the relative composition of clonal lineages 2A1, EC1, CO4, and US1 (*P. infestans*), and EC2 and EC3 (*P. andina*). Pie chart size is proportional to the total number of isolates per department, and numerical values indicate the number of isolates per clonal lineage.

Figure 4 illustrates the temporal distribution of *P. infestans* clonal lineages in Ecuador between 1993 and 2022. Throughout this period, EC-1 was the dominant lineage and was detected in most sampling years. In contrast, US1 occurred sporadically but was still detected in recent collections, including in 2022.

**Figure 4.**
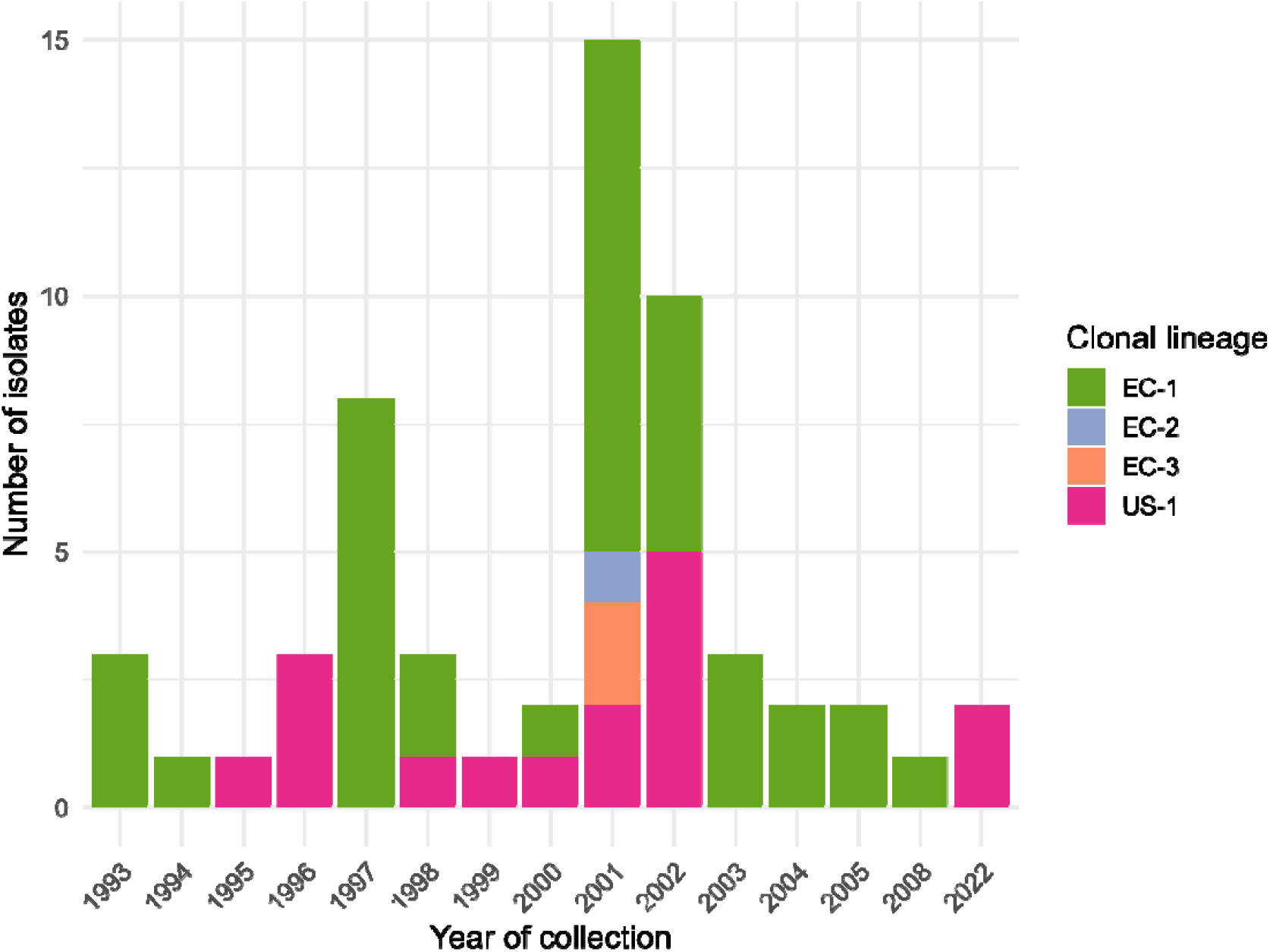
Clonal lineage distribution of *Phytophthora infestans* (EC-1, US-1) *and Phytophthora andina* (EC-2, EC-3) in Ecuador by year of collection (1993–2022)

#### Multilocus genotypes

Analysis of the 182 isolates yielded 126 multilocus genotypes (MLGs); including reference isolates, a total of 141 MLGs were detected (Table 2). The most frequent genotypes were MLG139 (n=22), MLG80 (n=7), and MLG131 (n=6) (Figure 5).

**Table 2.**
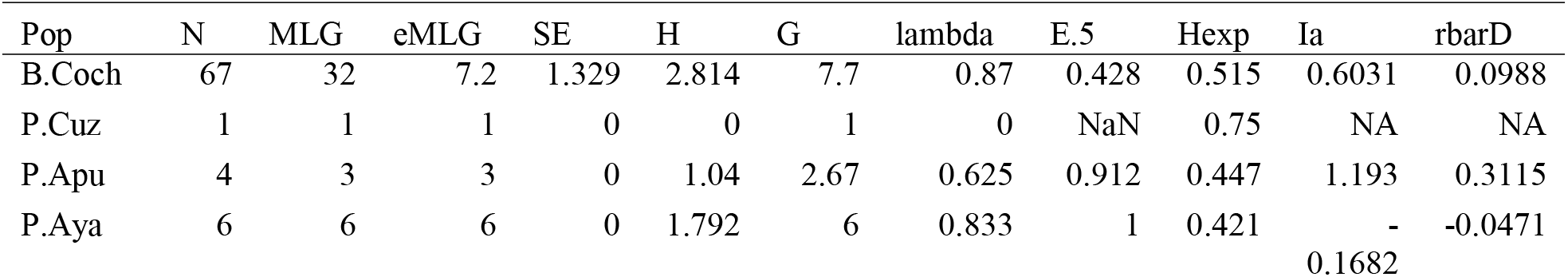
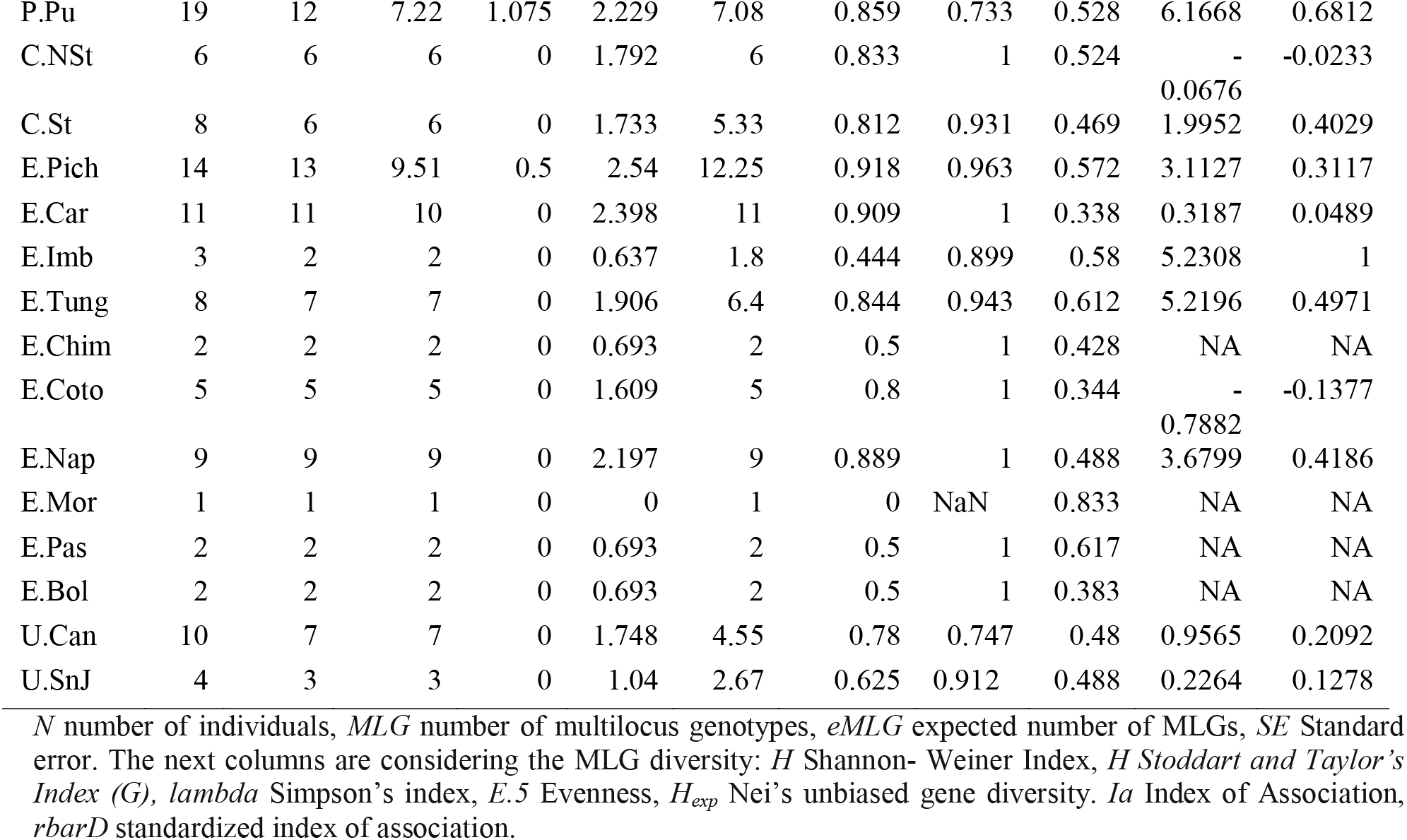
Diversity statistics for microsatellite data across 12 loci in *Phytophthora* populations from Bolivia, Peru, Colombia, Ecuador, and Uruguay categorized by location.

**Figure 5.**
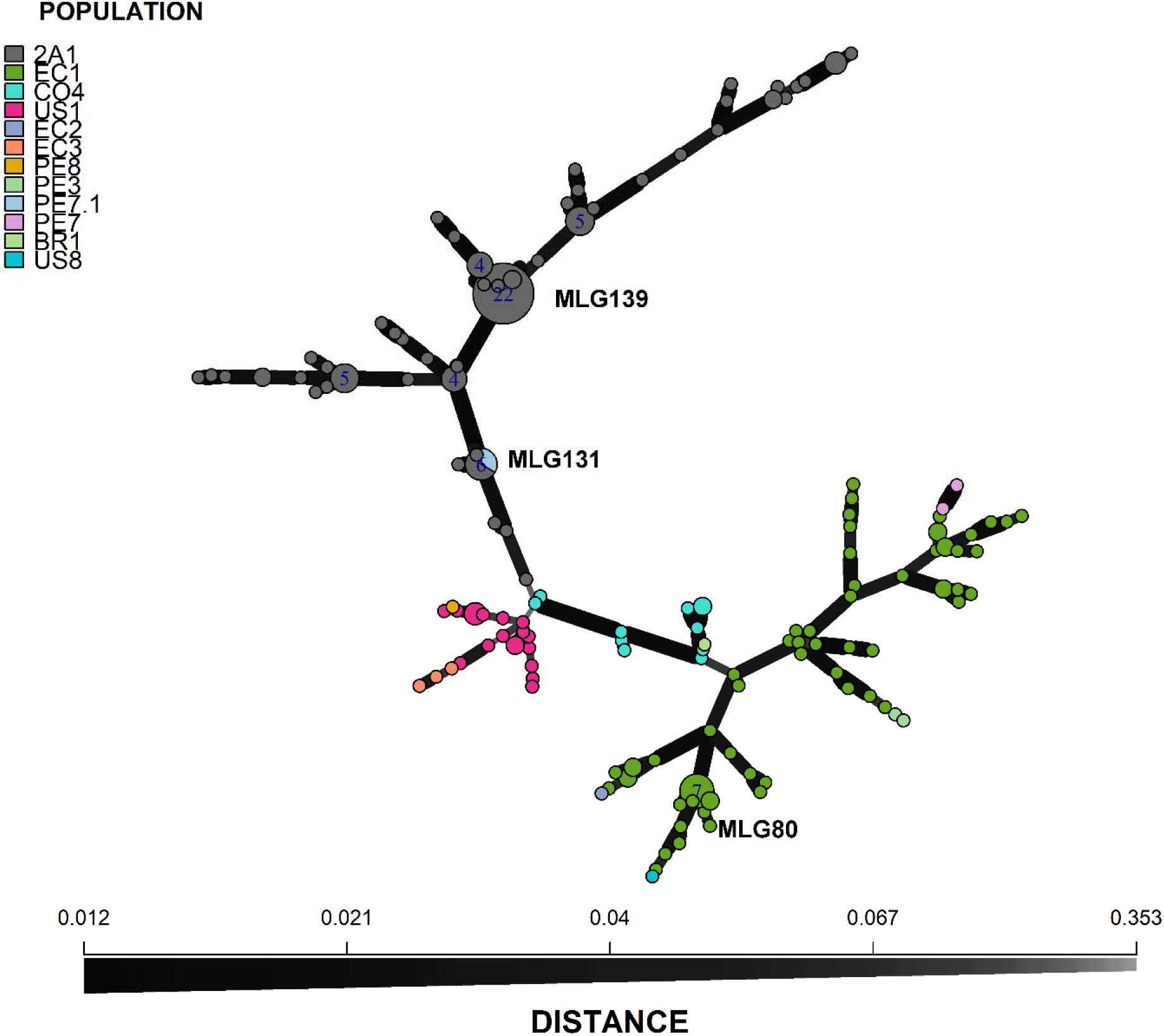
Minimum spanning network (MSN) of multilocus genotypes (MLGs) of *Phytophthora infestans* and *Phytophthora andina* populations from Bolivia, Peru, Colombia, Ecuador, and Uruguay. Each node represents a distinct MLG, with node size proportional to its frequency. Colors indicate clonal lineage assigned based on reference genotypes (US1, EC1, 2A1, PE3, PE7.1, PE7, BR1, US8 for *P*.*infestans;* EC2, EC3, and PE8 for *P*.*andina*). Edges correspond to Bruvo’s genetic distances between MLGs.

MLG80 corresponded to six isolates (one from Ayacucho and five from Puno) and one EC1 reference isolate, while MLG67 corresponded to one isolate from Puno and the second EC1 reference isolate. MLG131 included four isolates from Puno and the reference genotype previously designated PE7.1.

Puno showed relatively high genotypic diversity (MLG = 12, Hexp = 0.53) and strong signals of clonality D = 0.68). Similarly, in Ecuador, Tungurahua and Napo exhibited high genotypic richness (MLG = 7 and 9, respectively) together with elevated linkage disequilibrium indices (Ia = 5.22 and 3.68), indicating predominantly clonal population structure. Bolivia, which represented the largest number of isolates (n=67), showed substantial genotypic diversity with 32 MLGs, although the most frequent genotype (MLG139) corresponded to the clonal lineage 2A1 (Table 2, Figure 5).

D=0.37; p=0.001), confirming significant linkage disequilibrium.

#### Marker performance

The multilocus accumulation curve increased steadily with the addition of markers but did not reach complete saturation at 11 loci, indicating that each SSR marker contributed to genotype discrimination and that full marker coverage was necessary to capture the genetic diversity of isolates analyzed (Figure S3).

Among the SSR markers, G11 and D13 showed the highest numbers of alleles (18 and 17, respectively). The average of alleles per locus was 7.92, indicating moderate allelic diversity. Diversity indices (Simpson’s 1-D and expected heterozygosity, Hexp) ranged from 0.38 to 0.81, with mean values of 0.60. Evenness ranged from 0.47 to 0.89 (mean=0.72), indicating moderate variation in allele distribution across loci (Table 3).

**Table 3.** Population statistics for the 12 microsatellite loci in *Phytophthora* populations from Bolivia, Peru, Colombia, Ecuador, and Uruguay.

| locus | allele | 1-D<sup>b</sup> | Hexp<sup>b</sup> | Evenness<sup>b</sup> |
| --- | --- | --- | --- | --- |
| D13 | 17 | 0.67 | 0.67 | 0.47 |
| SSR8 | 5 | 0.64 | 0.64 | 0.84 |
| SSR4 | 12 | 0.81 | 0.81 | 0.8 |
| Pi04 | 4 | 0.61 | 0.61 | 0.89 |
| Pi70 | 3 | 0.4 | 0.4 | 0.72 |
| SSR6 | 7 | 0.54 | 0.54 | 0.77 |
| Pi63 | 3 | 0.61 | 0.61 | 0.88 |
| G11 | 18 | 0.7 | 0.7 | 0.53 |
| SSR3 | 7 | 0.61 | 0.61 | 0.67 |
| SSR11 | 5 | 0.62 | 0.62 | 0.8 |
| SSR2 | 3 | 0.38 | 0.38 | 0.63 |
| Pi4B | 11 | 0.59 | 0.59 | 0.63 |
| mean | 7.92 | 0.6 | 0.6 | 0.72 |

<sup>b</sup> 1-D, Simpson's diversity index. *Hexp*, Nei's gene diversity. Evenness indices of evenness.

#### AMOVA

AMOVA revealed significant genetic structuring among clonal lineages. A substantial proportion of the total molecular variance (57.77 %) was attributed to differences among clonal lineages, indicating strong genetic differentiation at this level. An additional 10.21% of the variance was explained by differences among geographic locations nested within clonal lineage. The remaining 32.02% of the variation occurred within samples. All hierarchical levels were statistically significant according to the Monte Carlo test (P = 0.001) (Table 4).

**Table 4.**
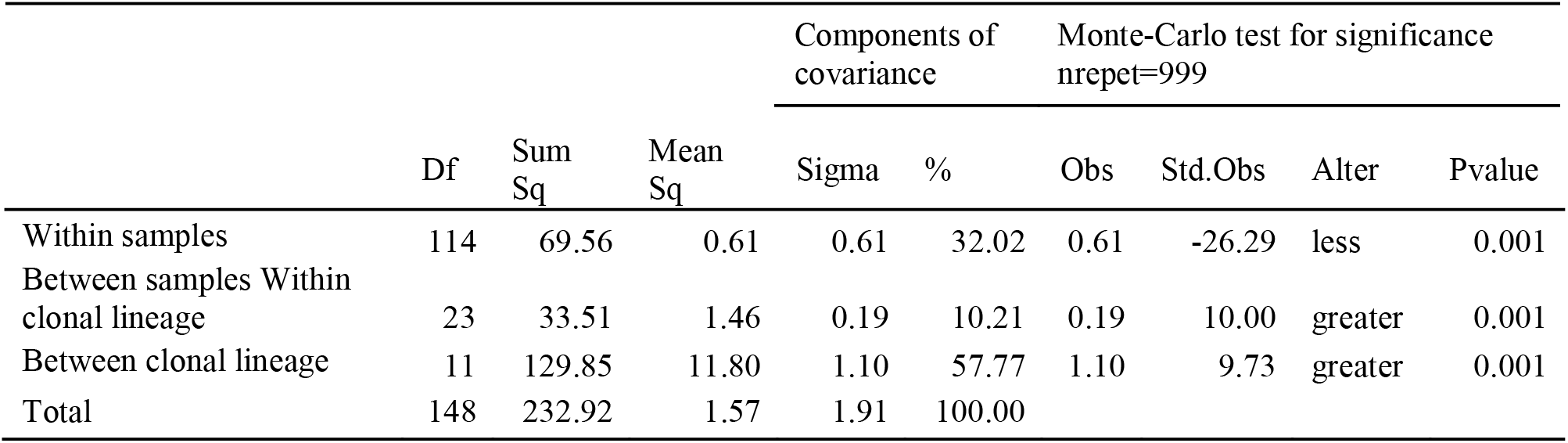
Analysis of molecular variance (AMOVA) of *Phytophthora* isolates from Bolivia, Peru, Colombia, Ecuador, and Uruguay.

## Discussion

The genetic composition of *Phytophthora infestans* populations is dynamic and continuously evolving, with important implications for late blight management. Previous studies have shown that *P. infestans* populations worldwide display substantial genetic diversity and undergo rapid evolutionary change (Cooke & Lees, 2004; Dey *et al*., 2018). Several studies have investigated *P. infestans* populations in the Andean region, where Peru has been proposed as a possible center of origin of the pathogen (Gómez-Alpizar *et al*., 2007; Patarroyo *et al*., 2024; Coomber *et al*., 2025).

Perez et al. (2001) analyzed *P. infestans* isolates from Cusco and Puno in Peru and identified EC1, PE3 and US1 as predominant clonal lineages infecting cultivated and wild potatoes (EC1 and US1) and cultivated potatoes (PE3). In the Peruvian highlands, host specificity does not appear to strongly influence the composition of pathogen populations infecting tuber-bearing Solanum species (Garry *et al*., 2005). More recent studies in these departments identified only EC1 and a new genotype named PE7.1 (Lindqvist□Kreuze *et al*., 2020). EC1 has remained the predominant genotype in Peru since its first report by Perez et al. (2001), whereas US1 and PE3 have not been detected in more recent population studies. PE3 was previously associated with native cultivated landraces and wild tuber-bearing species (Perez *et al*., 2001; Garry *et al*., 2005). Our analysis indicates that the genotype previously designated as PE7.1 does not represent an independent lineage but instead corresponds to a variant within the broader 2A1 clonal lineage, which has been reported in several South American countries (Acuña et al., 2019; F. Lucca et al., 2023). US1 was identified in Ecuador in both historical and recent collections, suggests that this lineage may persist at low frequencies in alternative hosts. Previous studies from the Andean region have reported *P. infestans* clonal lineages infecting non-potato *Solanum* species (Garry *et al*., 2005; Lindqvist□Kreuze *et al*., 2020), which may contribute to the persistence of pathogen populations outside cultivated potato fields. More broadly, *P. infestans* can infect a range of *Solanum* species, including wild hosts (Grünwald & Flier, 2005), and its long-distance dispersal is strongly associated with human-mediated movement of infected plant material, particularly seed tubers (Fry *et al*., 2015), allowing lineages such as US-1 to persist or re-emerge despite being largely displaced from potato populations. The long-term survival of clonal *P. infestans* lineages likely hinges on a fine balance between their aggressiveness, adaptability to environmental conditions, and capacity to counteract disease management strategies (Mariette *et al*., 2016; Fry, 2020). Our research also corroborated previous studies on the population structure of certain strains from Ecuador, which had identified the clonal lineages EC1 and US1(*P*.*infestans*), EC2 and EC3 (*P*.*andina*) —using RFLP fingerprinting —as being associated with different host species: EC1 (cultivated and wild potatoes), EC2 (wild potato), EC3 (cultivated tree tomato), and US1 (cultivated tomato species, pepino melon, naranjilla, wild potato species, wild relative of naranjilla) (Gómez-Alpizar *et al*., 2008; Oliva *et al*., 2010). Subsequent taxonomic studies suggested that isolates previously classified within the EC3 lineage may correspond to *Phytophthora betacei rather than P*.*andina* (Mideros *et al*., 2018). These findings were updated using the 12-SSR multiplex method. The identification of EC1 and the newly discovered CO4 in Colombia highlights a geographical influence: EC1 has been reported in Ecuador and Venezuela (Forbes *et al*., 1998; Garry *et al*., 2005), while the new clonal lineage CO4 was found only in the department of Norte de Santander, where a humid tropical ecosystem is present (Dueñas *et al*., 2007). Norte de Santander borders Venezuela, where the Ia mitochondrial haplotype and A1 mating type have been documented, which corresponds to the profile observed in our strains from Norte de Santander (Briceño *et al*., 2009; Cárdenas *et al*., 2011). We did not identify the US8 lineage, which was previously reported in Colombia and was isolated from Cape gooseberry and also possesses the Ia haplotype, but corresponded to the A2 mating type (Vargas *et al*., 2009).

The existence of both mating types promotes gene exchange between populations, as demonstrated by the discovery of distinct genotypes in fields where both mating types are present (Goodwin, S.B. *et al*., 1995). In 2001, Perez et al. (2001) found that the predominant mating type in Peru was A1, whereas in Bolivia, the A2 mating type was detected in the past (Goodwin, S.B. *et al*., 1994), which belongs to the clonal lineage BR1. The BR1 reference was added to our SSR analysis, and we determined that it belonged to a different cluster from the isolates collected from Bolivia in our study. Sexual reproduction may occur when compatible A1 and A2 mating types coexist in the same location; however, this was not observed in the populations sampled in this study. However, the risk of sexual reproduction is always present when both mating types co-exist, which is the case in potato production zones near the border between Peru and Bolivia.

*P. infestans* may have a broader host range than previously thought, which could have implications for disease management. In our study, the A1 mating type was detected using the W16 marker, which has been proven to be accurate in identifying mating types in clonal lineage EC1 (Brylińska *et al*., 2018). The pairing test on V8 agar was chosen for the Ecuadorian population because the W16 marker has shown inconsistent results for the US1 genotype and also for *P. andina*, as reported by Brylińska et al. (2018). In that study, all US-1 isolates of A1 mating type amplified the fragment typical of A2 using W16, making this marker unreliable for populations where these lineages occur.

The high proportion of genetic variation detected among clonal lineages (57.77%) indicates strong genetic differentiation among lineages. This pattern is consistent with strong genetic differentiation among clonal lineages identified using SSR markers and with the predominantly clonal population structure reported for *P*.*infestans* in many regions (Goodwin, S.B. *et al*., 1995). In contrast, the lower proportion of variance explained by geographic location suggests that lineage identity is a stronger determinant of genetic structure than regional distribution. Lineages EC1, EC2, and EC3 are predominantly associated with Andean countries, whereas lineage 2A1 has been reported mainly in Europe, while the origin of lineage US1 remains more controversial (Forbes *et al*., 1997; Mariette *et al*., 2016; Martin Md *et al*., 2016). These distinct evolutionary trajectories have likely generated strong genetic differentiation among lineages, which is reflected in the high variance component observed in the AMOVA. In a North American study of 77 isolates, Goodwin, S.B. et al. (1995) found that 63% was due to differentiation among lineages, and older lineages exhibited more pathogenic variation than recently introduced ones. The US1 clonal lineage has reportedly been replaced in Peru, with its presence limited to wild Solanaceae species in the 2016-2017 population. This change could be attributed to the lineage’s low virulence in cultivated potatoes (Lindqvist□Kreuze *et al*., 2020). Also, EC1 was reported to be adapted to higher temperatures than US1, and to exhibit a high level of resistance to metalaxyl (Garry *et al*., 2005), in contrast to US1, which showed over 50% sensitivity to the fungicide (Lindqvist□Kreuze *et al*., 2020). The dynamic nature of *P. infestans populations*, with the emergence and replacement of dominant clones over time, highlights the importance of continuous monitoring and studying the genetic variation in this pathogen for effective disease management strategies (Knaus *et al*., 2020). For example, in America, the US23 lineage, which is currently widespread in the United States, exhibits genetic similarities with the BR1 lineage found in Bolivia and Brazil, whereas other lineages in the U.S. are more closely related to populations in Mexico (Saville & Ristaino, 2019). In terms of management strategies, we need to consider the aggressiveness of clonal lineage EC1 that currently dominates in the southern regions of Peru (Lindqvist□Kreuze *et al*., 2020). Isolates within this lineage exhibit variation in key epidemiological components, particularly incubation period, infection frequency, and lesion expansion (RAULEC, a measure of disease progression over time)(Chacón *et al*., 2007), which may influence disease development and complicate management strategies. Furthermore, 2A1 shows low or moderate aggressiveness relative to the other coexisting lineages (Mariette *et al*., 2016). Additionally, the variations in aggressiveness observed in new populations might result from better adaptation to environmental conditions like temperature or humidity (Lindqvist□Kreuze *et al*., 2020).

*Phytophthora infestans* can be transmitted through infected seed potatoes, which represent an important source of primary inoculum (Kool & Evenhuis, 2023). The movement of infected seed tubers has been widely recognized as a major driver of long-distance dispersal of *P*.*infestans* lineages and late blight outbreaks worldwide (Guha Roy *et al*., 2021; Ryley & Drenth, 2024; Saffer *et al*., 2024; Rhouma *et al*., 2024).In several Andean production systems, the use of farm-saved seed potatoes and the exchange of planting material among neighboring countries may facilitate the persistence and regional spread of clonal lineages such as those identified in this study.

MLGs grouped together from the potato-growing areas of Puno EC1(MLG80) and 2A1 (MLG131) were found approximately 40 km from the Bolivian border. The current *P. infestans* population of Puno (EC1 and 2A1) is different from that of the old population (US1, PE3 and EC1) (Perez *et al*., 2001), suggesting that displacement of the old dominant lineages has occurred. These shifts in lineage composition, revealed by SSR genotyping, reflect temporal changes in population structure. The distribution of the 2A1 lineage observed in this study may reflect a broader regional pattern of introduction and spread. This lineage, originally reported in Europe and now dominant in several South American countries, may have been introduced into the Southern Cone (e.g., Argentina, Brazil, or Uruguay) through the importation of infected seed tubers, and subsequently expanded northwards into Bolivia. The presence of 2A1 in Bolivian populations and near the Peruvian border suggests a possible ongoing process of regional dissemination, potentially leading to its future establishment in Peruvian populations. In addition to regional connectivity, alternative introduction pathways cannot be excluded. The exclusive detection of the 2A1 clonal lineage in Bolivia suggests a relatively homogeneous population structure. While our results indicate possible connectivity with southern Peru, the origin and dissemination pathways of this lineage in Bolivia remain uncertain. Given the widespread global distribution of 2A1 and its association with international seed tuber movement, historical introductions through formal or informal seed systems cannot be excluded. The current population structure may therefore reflect a combination of regional spread and independent introduction events. Such dynamics highlight the importance of monitoring the spread of invasive clonal lineages across connected agricultural systems. Since 2011, a similar shift has been noted in Argentina for the oldest lineages, AR1-AR5_A2, and in Uruguay and Brazil for BR1_A2 (Forbes *et al*., 1998; Deahl *et al*., 2003), and all have been replaced by EU-2_A1 (2A1) (Lucca *et al*., 2019, 2023). The most abundant MLG (MLG139) was shared between seven provinces of Cochabamba-Bolivia, with 64.2% of MLGs being non-unique and distributed across multiple provinces, and the presence of clonal lineage 2A1, which is present in several South American countries, suggesting that it may be older and have had more time to spread. In the case of Peru, 56.7% of the MLGs were unique. The high number of unique MLGs identified using SSR markers highlights the substantial genetic variation within populations despite their predominantly clonal structure (Forbes *et al*., 1998; Dey *et al*., 2018). This difference in diversity could be attributed to the origin of the pathogen and its history of introduction (Saville *et al*., 2016).

Our findings demonstrate that *P. infestans* populations in South America exhibit strong lineage-based genetic structure shaped by regional ecological conditions and the movement of plant material. These results highlight the need for transnational genetic surveillance strategies to monitor the emergence and spread of clonal lineages that may affect late blight management.

## Author contributions

MI: coordination, sample processing, molecular laboratory work, marker scoring, analysis, interpretation of data, and drafting of the manuscript.

MC: sampling

WP: coordination, interpretation of data and revision of the manuscript

LS: sampling, sample processing

SG: sample processing.

DV: sampling

VC: coordination

BG: coordination, interpretation of data

JK: interpretation of data and revision of the manuscript.

## Acknowledgements

We would like to thank Freddy Ventura and Marcelo Vinueza for their technical support during the sample processing.

## Data availability statement

Not applicable

## Ethic declaration

Not applicable

## Supplementary Information

Supplementary materials are provided as separate files.

