## Supplementary material for "Population structure of *Phytophthora infestans* collected from potatoes in Ecuador, Colombia, Peru, Bolivia and Uruguay": Figure S1, Figure S2, Figure S3

**Supplementary Information**

Supplementary Table Legend

**Table S1**. Metadata for the *Phytophthora infestans* isolates used in this study, including geographic origin, host, year of collection, and genotyping data. The column “source” indicates previously published isolates used as references, with the corresponding citations listed in the bibliography.

Supplementary Figures

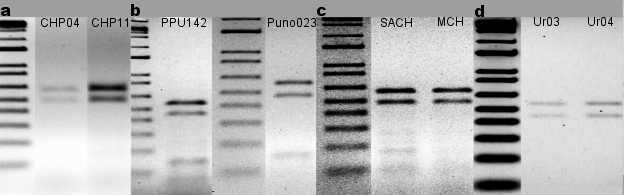

**Figure S1**. HaeIII Restriction Enzyme Digestion of PCR using W16 Primer Set in Selected a. Bolivian (CHP04, CHP11), b. Peruvian (PPU142, Puno023), c. Colombian (SACH, MCH), and d. Uruguayans (Ur03, Ur04) strains. Ladder 1kbplus.

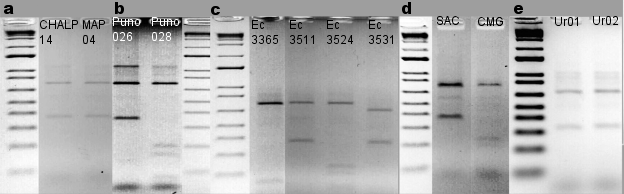

**Figure S2**. Results of Restriction fragments from digestion of the P2 region with enzyme *MspI* with **a**. Bolivian (CHALP14, MAP04), **b**. Peruvian (Puno026, Puno028), **c**. Ecuadorian (Ec-3365, Ec-3511, Ec-3524, Ec-3531), **d**. Colombian (SAC, CMG), and **e**. Uruguayans (Ur.1, Ur.02) strains. Ladder 1kbplus.

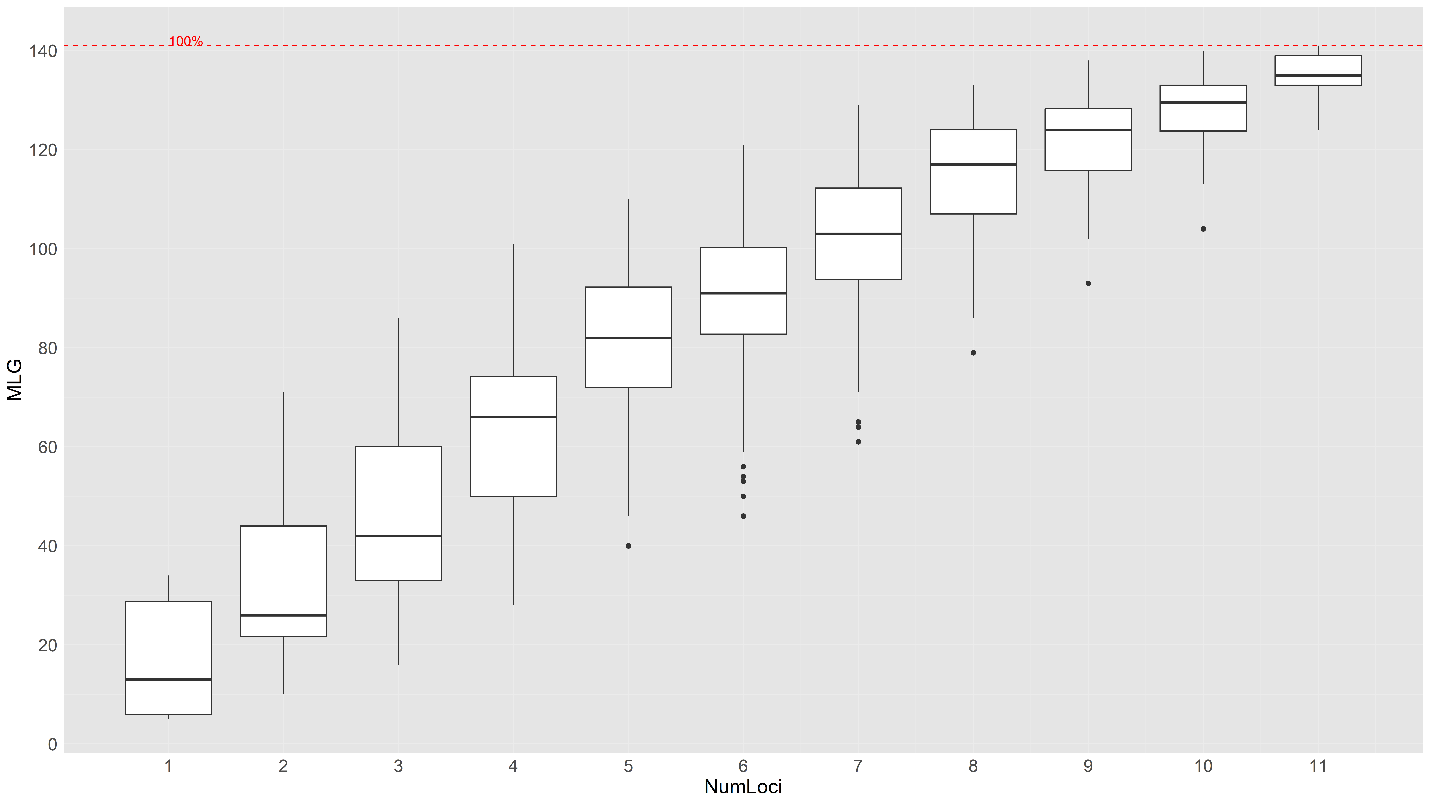

**Figure S3**. The genotype accumulation curve. Numloci refers to the number of SSR loci, whereas MLG denotes the number of multi-locus genotypes. The MLG resolution was depicted as 100%, as indicated by the dashed red line.
